# Rapid on-site universal vertebrate species identification via multi-barcode nanopore sequencing

**DOI:** 10.1101/2025.06.04.657926

**Authors:** Emily C. Patterson, Sandrella Morrison-Lanjouw, Mark A. Jobling, Jon H. Wetton

## Abstract

The growing illegal wildlife trade (IWT) threatens biodiversity and is a conduit for zoonotic disease, yet its risk of detection is low. Once processed, trafficked species are difficult to identify morphologically, and currently require DNA-based approaches that are time-consuming, costly, and lab-based. There is thus a need for a rapid, cheap, on-site method for species identification. We describe VeRIF-ID (Vertebrate Rapid In-Field Identification via DNA), a method that employs simultaneous on-site nanopore sequencing of four different mitochondrial DNA barcodes. Primers were designed to produce short amplicons to aid analysis of damaged DNA, and to be effective over a broad taxonomic range of vertebrates from lamprey to chimpanzee. Validation demonstrated species-level identification in 91% of 83 tested species, and genus/tribe-level identification of the remaining species (which are also problematic with existing approaches). DNA extraction, PCR and library preparation steps were simplified and optimised so that sampling to species identification takes <3 h for a single sample. Species components are identifiable non-quantitatively in prepared mixtures of muscle tissue from up to five species, and laboratory tests of Traditional East Asian Medicine samples reveal DNA from species including critically endangered saiga antelope and black rhinoceros. In conjunction with a portable BentoLab device the necessary equipment and reagents are easily portable, and we apply the method to analyse seized bushmeat and fish samples within an airport customs zone, identifying mammal and fish species in 15 samples within 6 h. The initial equipment costs for VeRIF-ID are ∼$8K, and the cost per sample of ∼$10-48 (depending on set-up), considerably cheaper than current conventional lab-based approaches. The method requires only basic hands-on skills. Ongoing trials with potential end-users will focus on establishing forensic reporting criteria prior to casework implementation. Future development of user-friendly bioinformatic interfaces will aim to fully democratise species identification.

## Introduction

The trading of wildlife for food (bushmeat), traditional medicinal ingredients, and as pets, involves approximately a quarter of all terrestrial vertebrate species globally [1]. The illegal wildlife trade (IWT) is one of the most lucrative illicit industries, with an estimated worth of $7 - 23 billion per year, and a growth of between 5 - 7% annually [2]. Both the probability of detection and the penalties, when detected, are generally low [3]. The deleterious consequences include extinction, biodiversity loss, the introduction of invasive species, and the emergence of novel zoonotic diseases [4]. The trade poses a threat to all 17 United Nations Sustainable Development Goals (SDGs), and most clearly in SDG 16 (Peace, Justice and Strong Institutions) and SDG 8 (Decent Work and Economic Growth) [5]. Despite this, eliminating the IWT has only recently been recognised as a political priority [6, 7].

The detection and seizure of illegal wildlife in the supply chain, for instance at border posts, ports, and airports, represents an essential measure for the enforcement of trade regulations [8]. Animal carcasses can be recognised through visual inspection, and their identification may be aided by the use of visual guides (for example [9]). However, processed wildlife products lacking discernible morphological characteristics (e.g. meats, fish fillets, animal parts and derivatives within products such as Traditional East Asian Medicines [TEAMs] [10]) are frequently encountered. Processing is often carried out to conceal the species of origin and can make products visually indistinguishable from those obtained from farmed or legally harvested species [11]. For example, 59% of 194 items of meat seized at Brussels Zaventem airport were unidentifiable morphologically [12].

Species substitutions are of particular concern in fisheries. Each year, illegal, unreported, and unregulated (IUU) fishing is thought to cost about $36.4 billion in losses [13]. Widespread food fraud throughout the fisheries logistics network has seen high-value species adulterated or substituted with low-value species, and imported seafood labelled as domestic [14]. Equally, illegally harvested protected species may be sold as low-value species [15], as seen, for example, in the UK where protected shark species have been a component of traditional fish and chips [16]. Further food fraud and adulteration cases include the presence of pork in seafood products [17], or horse-meat in beef products [18], causing serious ethical or religious issues for consumers.

Species identification in processed samples depends on DNA-based analysis, which is normally achieved through DNA barcoding [19], the analysis of sequences that display low intraspecific but high interspecific variation. DNA barcodes used for animals lie in the mitochondrial genome (mtDNA): a segment of ∼600 bp in the cytochrome c oxidase subunit I gene (COI), is the principal barcode [19, 20], while other commonly used barcodes include sequences in cytochrome b (CYTB) [21–23], 16S [24, 25] and 12S ribosomal RNA (rRNA) genes [26, 27]. In the barcoding process, DNA is extracted from the sample and the chosen barcode is amplified by PCR using conserved primers.

Typically, such primers are not applied across multiple taxonomic groups, but are targeted at a particular group, for instance shark-specific primers for use in pet foods [28], or primers designed for the identification of tortoises [29], or crocodiles [30]. Amplicons are usually then Sanger-sequenced and the resulting sequences compared to a relevant database (e.g. the NCBI nucleotide database, or the Barcode of Life Data System (BOLD) [31]) to identify the best-matching species.

This approach has a number of disadvantages. Time-consuming and costly research and development is needed every time a new taxon-specific species identification system is developed. Testing is done in a laboratory with expensive equipment, specialist staff, and associated costs, so that each test costs >$300 {PAW Forensic Working Group, 2024 #4993}. The infrastructure requirements mean that samples seized at customs posts or in the field must be transported to a laboratory. This complicates the evidential chain, incurs additional time and expense, introduces biosecurity risks, and can add the administrative burden of satisfying CITES (Convention on International Trade in Endangered Species of Wild Fauna and Flora) restrictions if export and import across national boundaries are necessary. There are also potential issues with identification via the barcoding process itself. Sequence databases are not free of errors [32, 33], and mislabelling of sequences or the erroneous database inclusion of sequences of nuclear mitochondrial segments (numts; [34, 35]) can provide inaccurate identifications [36]. This is more likely to occur when only a single barcode, e.g. COI, is used in a test [33]. Use of single barcodes also risks misidentification due to the co-amplification (or preferential amplification) of numts in samples undergoing testing.

A number of mitigations can be envisaged for these problems. ‘Universal’ PCR primers that amplify barcode sequences across a very wide taxonomic range could provide an agnostic test allowing a single test to be applied to any suspicious item whilst also reducing validation, training and consumables costs. Simultaneous amplification and sequencing of multiple barcodes could mitigate against problems with databases and numt amplification. Finally, on-site analysis could produce results more quickly, reinforce the chain of custody, and facilitate the application of penalties on those illegally trading in wildlife products. The need for portable tools to carry out rapid on-site species identification has been emphasised by recent reports and surveys [37, 38].

The portability, low start-up costs and ready availability of the Oxford Nanopore Technologies (ONT) MinION sequencing device make it a good candidate for in-field species identification via DNA barcoding. Indeed, it satisfies the key cost and accuracy requirements of a field-based system for wildlife forensic investigations as determined by surveying wildlife forensic and law enforcement professionals [37]. The MinION has an initial cost of $2999, weighs just ∼100 g, and is plugged directly into a laptop USB port for operation and sequence interpretation. This device has already been used to identify species, from diverse vertebrates in Tanzania [39], to reptiles and amphibians in the Ecuadorian Chocó rainforest [40], a shark fin specimen from a market in India [41], as well as closely related plant species [42], and non-invasively collected wildlife samples [43]. Sequencing of a 421-bp CYTB segment in vertebrates has shown that ONT MinION data in conjunction with appropriate bioinformatic tools can generate sufficiently accurate consensus DNA sequences for forensic genetic species identification [44]. Protocols have been developed for species identification that, depending on sample type, can be completed in 10 h by a skilled operator in a laboratory [45].

In this study, we design, validate, and test for on-site application a system that addresses many of the problems associated with traditional DNA barcode-based approaches. Our system, VeRIF-ID (Vertebrate Rapid In-Field Identification via DNA), uses nanopore sequencing of four short DNA barcodes amplified with ‘universal’ primers that are diagnostic over a very broad taxonomic range. We simplify and optimise DNA extraction, PCR and library preparation such that the process from sampling to species identification takes less than 3 h for a single sample and 6 h for a batch of 24. In conjunction with a portable BentoLab device, the method is suitable for on-site application, as demonstrated by bushmeat species identification within a major airport’s customs zone.

## Materials & methods

### Primer design

Whole mitogenome sequences for a phylogenetically diverse range of mammal (n = 113), fish (n = 59), bird (n = 35), reptile (n = 27), and amphibian (n = 15) species were downloaded from GenBank (Table S1). Up to 20 sequences per species were included. The FeatureExtract 1.2 Server (https://services.healthtech.dtu.dk/service.php?FeatureExtract-1.2) was used to extract COI, CYTB, and 16S and 12S rRNA gene sequences. A multiple sequence alignment of the resulting 843 sequences was carried out using Clustal Omega [46] on the European Bioinformatics server. JalView [47] was used to inspect the resulting alignment and barcode primers were designed such that their 3’ ends were in conserved blocks bracketing variable regions. For each barcode, the variable region between primer sites was extracted and used as input with MEGA (v10.1.8) to build a neighbour-joining (NJ) Kimura two-parameter (K2P) tree with 100 bootstrap replicates [48]. The tree was examined to visualise the discrimination power of the primer pair under consideration. Primers favouring shorter amplicons were preferred given the known failure of longer barcodes in more degraded DNA samples [49, 50]. Primers were designed to give amplicons in the length range 150 – 353 bp (Table S2).

### Sample collection

Single-source samples encompassed a wide range of species and sample types, and are listed in Table S3. A further set of samples of muscle tissue from seven species (cattle, sheep, pig, chicken, turkey, salmon, and trout) were used to create lab-prepared mixtures. Finally, Traditional East Asian Medicine (TEAM) samples confiscated by the Metropolitan Police Service (London, UK) were analysed. Tissue, blood, and DNA extracts were stored at -20°C with all other sample types kept at room temperature prior to DNA extraction.

### DNA extraction

Hair, shed reptile skin, hedgehog quill, frog spawn, claw, eggshell, and feather samples were washed twice in 500 µL 70% (v/v) ethanol and twice in 1 mL Milli-Q water. DNA was extracted from 1-2 mm^3^ of claw, tissue, shed skin, feather calamus, spawn, quill, 3 µL of blood, or 5 – 10 hairs. A previously developed alkaline extraction method [51] was used for DNA extractions. Samples were heated in 100 µL of lysis solution (200 mM KOH, 2 mM EDTA, 0.2% (v/v) Triton X-100) at 98°C for 10 mins and then neutralised with 300 µL of 100 mM Tris-HCL. This extraction method was used with 0.01 g of each of the seven tissue samples used to create 19 lab-prepared mixtures (Table S4). Bone and scute samples were wiped down successively in bleach, water (x 2), and ethanol. After drying for 5 – 10 min, a hand-held drill (Ozito Cordless drill driver) with a 6-mm wood drill-bit was used to obtain ∼200 mg of bone powder. About 200 mg of pill, paste, or oil were obtained from TEAM samples (Table S5) for DNA extraction. Pills were powdered using a mortar and pestle. Samples were incubated at 56°C for 2 h with 950 µL 0.5M EDTA, 50 µl 1% (w/v) n-lauroylsarcosine sodium salt, 14 µL of 1M DL-DTT, and 32 µL proteinase K (20 mg/ml) [52]. After centrifugation at 11,300 g for 3 min, three 125 µL fractions of the supernatant were concentrated using the MinElute PCR Purification kit (QIAGEN) with the DNA eluted in 30-60 µL of buffer.

The QIAamp DNA Investigator kit was used for DNA extractions from ∼0.5 cm² of each TEAM plaster (Table S5) following the manufacturer’s protocol for the isolation of total DNA from paper and similar materials. DNA was eluted in 30-60 µL of water.

### PCR/Tetraplex amplification

HPLC-purified primers with an additional tandemly repeated 13-bp sequence at the 5’ end of each of the four barcode primer pairs (CYTB: [AGTGTCCTGCTAG]_2_; COI: [CTGAGGTGATCAG]_2_; 16S:[AACATGTGGTAAG]_2_; 12S: [TTATTCGCACACT]_2_ [53]) were used during amplification (Table S2).

These extra nucleotides were included to enable the addition of transposome-introduced indices at the 5’ end of primer sequences. Each reaction was performed in a 25-µL volume containing 12.5 µL of Type-It Master Mix (QIAGEN), 0.76 µL of forward and reverse CYTB primers (0.3 µM final concentration), 0.62 µL of forward and reverse COI and 16S primers (0.25 µM final concentration), and 0.2 µL of forward and reverse 12S primers (0.08 µM final concentration), plus 2-3 µL DNA extract. Cycling conditions consisted of an initial denaturation at 95°C for 5 min, then 35 cycles at 95°C for 30 s, 45°C for 90 s, 72°C for 30 s, and a final extension at 68°C for 10 min.

### Nanopore flowcell runs

A total of 96 single-source samples were prepared for sequencing on a MinION R9.4.1 flow cell (FLO-MIN106D, ONT). Mixtures and a no-template control were sequenced on a second flow cell. A 14-µL volume of PCR products was purified using 1.8× AppMag PCR Clean Up Beads (Appleton Woods Ltd), and the DNA eluted in 14 µL of water. The Rapid Barcoding kit 96 (SQK-RBK110.96, ONT) was used to prepare samples for sequencing where the manufacturer’s protocol was followed with some exceptions. The name of this kit uses the term ‘barcoding’ to describe the inclusion of short sequences (known as barcodes, or indices) in library preparation that allow data from multiple samples to be deconvoluted into individual sequence datasets. We refer to this process hereafter as ‘indexing’ to avoid confusion with barcoding in the sense of species identification. Sample quantification and normalisation steps were omitted. Instead, a 9-µL volume of each sample was used during the fragmentation and nanopore index addition step (‘tagmentation’), and a 9-µL volume of each tagmented sample was pooled together. Half of this volume was used during sequencing of the single-source samples, whereas the full volume was used for sequencing of the pooled mixed samples. Sequencing runs were carried out on a MinION sequencer and the software program MinKNOW (v21.06.0 for the single-source samples and v21.11.7 for the mixtures).

### Data analysis

Guppy (v5.0.16) (https://nanoporetech.com/software/other/guppy/history?version=5-0-16) was used for basecalling with the high accuracy model, and demultiplexing with the trimming index option enabled. The resulting sequences were filtered using NanoFilt (v2.8.0) [54] to select for sequences with a quality score of ≥11 and length between 100 and 500 bp. Reads assigned to a given nanopore index were processed using Porechop (v0.2.4) [55] to search for each of the forward and reverse barcode primer sequences and assign reads to their respective species ID barcode. The NGSpeciesID pipeline (v0.1.1.1) [56] was used to generate consensus sequences from the resulting groups of reads. A cluster abundance ratio of 0.1% was applied for the mixed flow cell run. Consensus sequences were used as a BLAST query against a local NCBI nucleotide database (constructed 02/02/2022), with the output file format specified via the option -outfmt “7 qseqid qlen evalue qcovs pident staxids stitle scomnames”. The BLAST output file, sorted by E-value, was then sorted by query sequence ID and percent identity before filtering to remove duplicate lines if the same taxonomic identifier was present more than once for any given sequence, and then further filtered to retain only the top hit(s).

In the single-source samples sequencing run, consensus sequences were discarded if they had <100 supporting reads, if the similarity for a given top match was <90%, or the BLAST hits were not labelled to the species level, or were of non-coding RNA sequences. In the mixed samples sequencing run, consensus sequences were retained if they had >20 supporting reads. This cut-off, which is about three times the highest number of supporting reads seen in the negative control, was selected to overcome any instances of contamination. A species was only considered present if forward and reverse consensus sequences were generated for at least two species ID barcodes in the mixtures run. In the single-source sequencing run, results for a species ID barcode were considered only when consensus sequences were generated in both the forward and reverse direction (though see Supplementary text).

In selected cases, all reads assigned to a nanopore index were aligned to a mitochondrial reference sequence using minimap2 (v2.24) [57]. The SAM alignment file was transformed using SAMtools (v1.13) [58] and visualised using IGV (v2.3.82) [59].

### On-site species identification

The methods used for on-site species identification at Twycross Zoo and Brussels Zaventem Airport varied depending on the sample source and site. Details are provided in Table S6.

## Results

We set out to design a rapid, robust and cost-effective system for vertebrate species identification that could be applied on-site with its only external requirement being a power supply. Below we describe the development and validation of this system, VeRIF-ID (Vertebrate Rapid In-Field Identification via DNA), and also give proof-of-principle examples of applications on-site, including deployment in a customs reception site within a major European airport.

### Primer design and in silico assessment of barcode performance

We first designed a new set of ‘universal’ primers for the amplification of four commonly used barcodes (COI, CYTB, 12S and 16S rRNA) for species identification in vertebrates. Design of previously published systems was based on a limited number of sequences and species, and given the growing availability of sequence data we began afresh. We used 843 sequences from a phylogenetically diverse range of 249 mammal, bird, fish, reptile, and amphibian species for primer design, aiming for short amplicons to maximise success in analysing degraded DNA. This resulted in novel 16S rRNA and COI primers that are quite distinct from other published primers; however, for CYTB and 12S rRNA, similarities to existing primers are notable (Figure S1), indicating that the inclusion of additional sequences made little difference here. The discrimination power of the four new primer pairs, which generate amplicon lengths between 150 and 353 bp, was assessed by building NJ K2P trees and seeking instances where a species-level identification was not possible (Figures S2-5). Across all barcodes, an incorrect species-level identification was seen in only two cases; for a tolai hare (*Lepus tolai*) and a saltwater crocodile (*Crocodylus porosus*). These results may be explicable by introgression previously noted in hares [60] or by sequence mislabelling in GenBank [61] and/or hybridisation in the crocodile. Species-level identification was achieved across the remaining 99.2% (247/249) species by examining all four barcodes.

### Development of a simple and rapid workflow for VeRIF-ID

The adoption of portable on-site species identification is dependent on the development of a fast, affordable, accurate and user-friendly system [37]. We examined each stage of the DNA barcoding process with these considerations in mind. Although commercial DNA extraction kits have been favoured in MinION-based barcoding studies [40, 62–64] because they yield extracts of high quality and concentration, these methods require several items of equipment, involve multiple manipulation steps, and have a high cost per sample (∼$4). Instead, we assessed previously developed DNA extraction methods that use commonly available laboratory chemicals [51, 65]. We settled on a two-step method [51] based on an alkaline (KOH) lysis buffer. Incubating samples for 10 min at 98°C in this buffer, followed by neutralisation, produced DNA extracts of suitable quality for downstream barcoding steps without tube-to-tube transfers. This ∼10-min extraction method worked well across a wide range of sample types (blood, muscle tissue, skin, feathers, hair, and claws [66]) and reduced the extraction time from ∼30 mins for blood, ∼1-3 h for tissue and hair (using commercial kit-based extractions) or 5 - 12 h for feathers [67], and over 24 h for claws [68]. A second extraction method was necessary for bone and Traditional East Asian Medicines (TEAM) samples, and a ∼2.5-h protocol was chosen based on a previously developed method [52] versus >12 h using a DNA extraction kit. These simpler extraction methods considerably reduced the cost of extraction per sample from ∼$7 using a commercial kit to <$0.10 using a KOH-based DNA extraction method, or ∼$5 using the EDTA-rich DNA extraction method for bone and TEAM samples.

To maximise efficiency, we set up a PCR tetraplex to co-amplify the four barcodes. The redundancy afforded by multiple barcodes favoured this approach despite the expected potential loss of barcode sequence data due to competition between targets. Heating steps accounted for ∼45-47% (∼47/100 or 54.5/120 min) of the time taken for PCR during on-site tests.

Oxford Nanopore Technologies’ rapid indexing kit (see Materials & Methods for an explanation for the name we use here) was selected to prepare samples for sequencing. The rapid kit was chosen despite manufacturer recommendations to use it with longer (500 bp - 5 kb) fragments of DNA (https://community.nanoporetech.com/docs/prepare/library_prep_protocols/rapid-sequencing-v14-amplicon-sequencing-sqk-rbk114-24-or-sqk/v/raa_9198_v114_revh_29nov2023), and its relatively low throughput compared to the ligation kit. The choice is for speed and simplicity: the method comprises just two main steps, takes only 10 minutes, and requires no third-party reagents. The short turnaround time is largely due to the use of a transposase complex that fragments DNA and simultaneously attaches nanopore indices to both newly-created 5’ strand termini (https://store.nanoporetech.com/rapid-barcoding-sequencing-kit-24-v14.html) during the first main step (‘tagmentation’). This simplifies preparation for sequencing by significantly reducing the number of pipetting and tube transfer steps (by 82 and 72% respectively compared to the native indexing kit (SQK-NBD114.24)) without sacrificing sequencing accuracy. The use of the rapid indexing kit does introduce some complexities in bioinformatic processing of generated sequence data which are discussed in Supplementary Text (see also Figure S6).

Samples were sequenced on a flow cell connected to a laptop with an offline version of the operating software MinKNOW and a local copy of the nucleotide NCBI database, negating the need for an internet connection. The workflow summarising these steps as well as the equipment and reagents required, which together fit into a standard luggage item, is shown in Figure 1.

**Figure 1:**
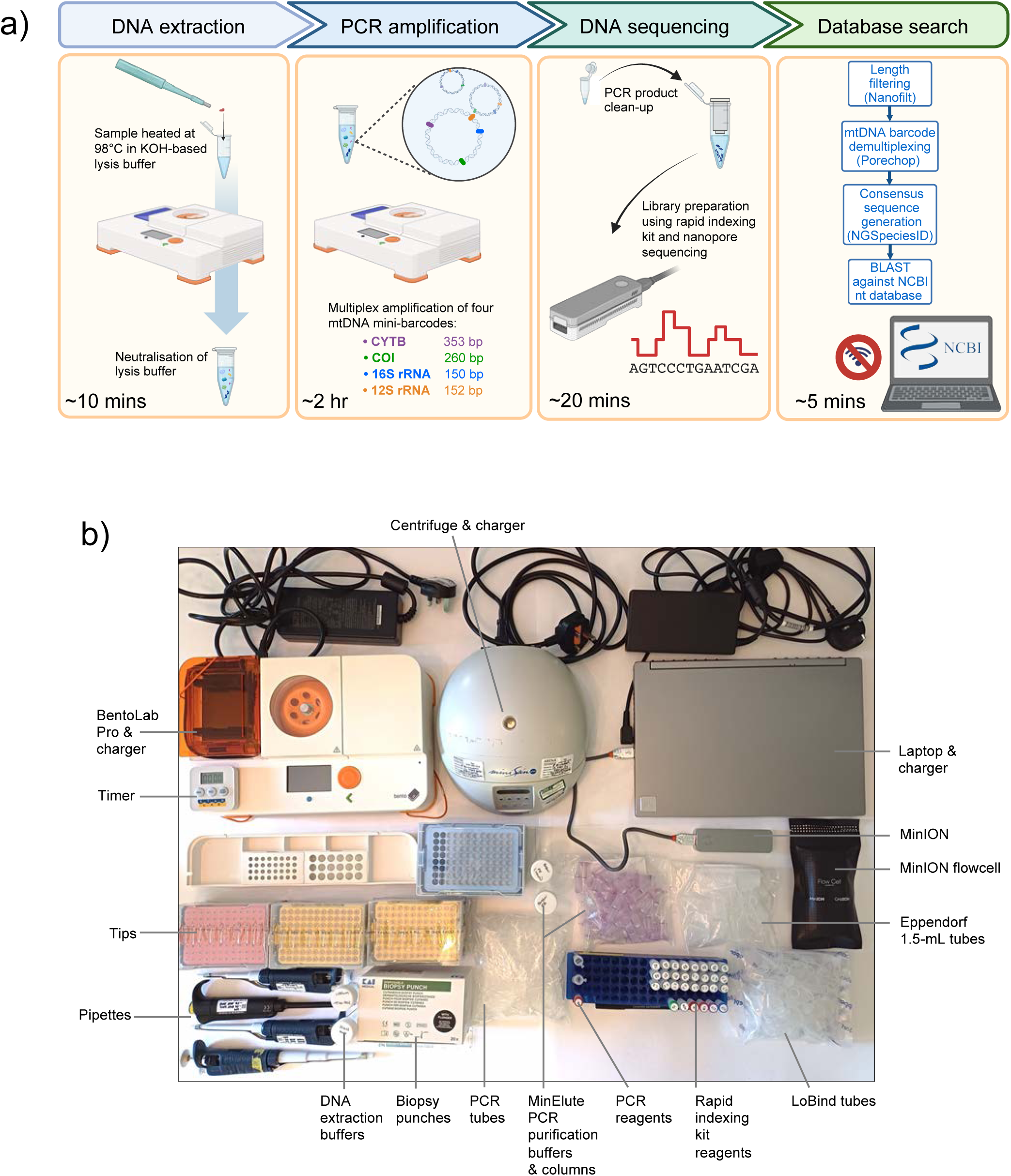
VeRIF-ID workflow and equipment. a) Schematic workflow for species identification. b) Photograph showing the necessary equipment and reagents, as deployed in Brussels Zaventem airport. See also Supplementary Text.

### Testing VeRIF-ID on single-source samples

We first tested the developed method on a nanopore flow cell to identify the species of origin for 96 single-source samples, originating from 45 mammals, 19 birds, 12 fish, six reptiles, one amphibian, and two invertebrates (Table S3). A total of 12.1 million reads were obtained (for details of Nanopore run statistics see Table S7), of which 88% were assigned to a nanopore index. An average of 58K reads per sample (range, 10K - 164K) were taken forward. Because of the mode of action of the rapid indexing kit used for library preparation, it was not possible to obtain a single consensus sequence per species ID barcode. Instead, forward and reverse consensus sequences were obtained for each species ID barcode (see Supplementary Text). Despite the transposase-introduced sequence resulting in longer than expected consensus sequences, and the use of a fragmentation-based kit, the alignment lengths between the query barcode consensus sequence and top database sequence match were generally in agreement with expected barcode insert sizes (Supplementary Text).

Of the 83 unique vertebrate species (Table S3), 75 could be assigned to species level, seven to genus, and one to tribe level (Figure 2). The eight less precisely assigned species include recently diverged domesticates, and are detailed in Table S8. For a further seven species (African pygmy hedgehog [*Atelerix albiventris*], Argentine anchovy [*Engraulis anchoita*], Malagasy giant rat [*Hypogeomys antimena*], great grey owl [*Strix nebulosa*], red crested turaco [*Tauraco erythrolophus*], tope shark [*Galeorhinus galeus*], and pink-backed pelican [*Pelecanus rufescens*]) there were missing or partial reference data for some barcode sequences (Table S8). This meant that alternative candidate species appeared as the top match(es) in the absence of data from the true species. Unexpected results were obtained in a further three instances, due to a range of issues (Table S8). Distinguishing nyala [*Tragelaphus angasii*] from bushbuck [*Tragelaphus scriptus*] is affected by NCBI reference sequence mislabelling [69]. Unexpected species assignment was also seen for the herring (*Clupea harengus*) and Arctic cod (*Boreogadus saida*) samples, which were identified as saithe (*Pollachius virens*) and Atlantic cod (*Gadus morhu*a) respectively. These results appeared genuine based on a reference-based approach where reads were aligned to the expected and assigned species. This surprising outcome is consistent with the finding that 55% of shop-bought fish samples tested in the UK were mislabelled [70].

**Figure 2:**
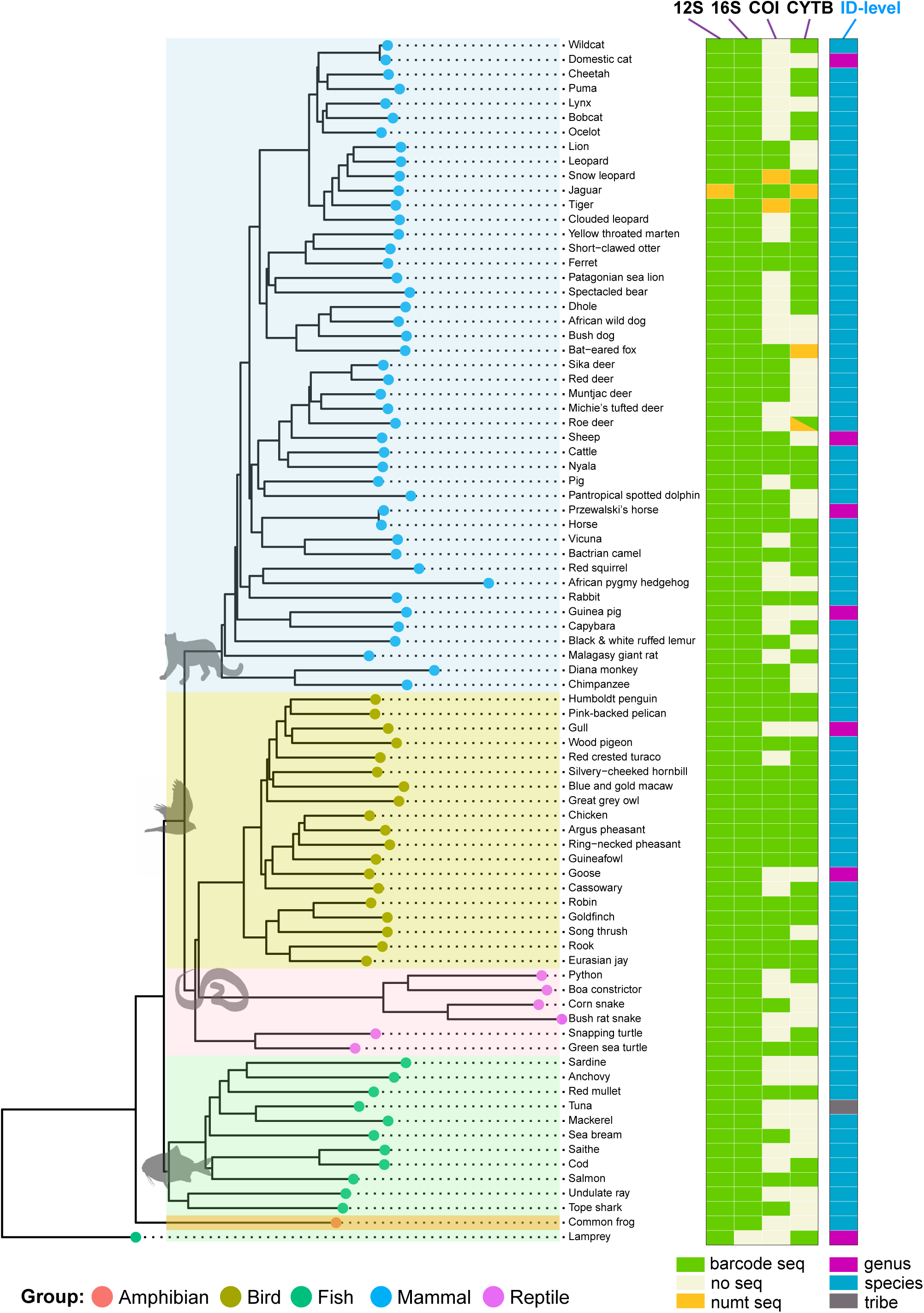
Barcode recovery across 83 vertebrate species following a nanopore flow cell run. To the left is a NJ K2P tree, created using publicly available cytochrome b sequences from GenBank for the species (or closely related species) displayed. Recovery of barcode and numt sequences for each of the four tested barcodes is shown schematically to the right, as well as the taxonomic level of identification achieved. Sequences were labelled as numts if their top match was to nuclear DNA and/or the top mitochondrial match was in disagreement with the other barcode sequence matches. Two additional invertebrate species were also analysed with some success [66] but are not shown here.

### Species identification in mixtures

Because tested samples often contain mixtures of material from more than one species, we carried out a qualitative assessment of species identification in known mixtures of vertebrate muscle tissues. Nineteen lab-prepared mixed samples (Table S4) and one negative amplification control were sequenced on a flow cell. A total of 2.5 million reads were obtained, and of these, 1.4 million remained after filtering, and 988K (70%) were assigned to their respective species ID barcode. It was possible to successfully identify all expected species in mixtures with up to five species of mammals and birds. It is important to note that here we consider only the presence or absence of a species, rather than a quantitative assessment. As shown schematically in Figure S7, fewer reads than expected are observed for some species in particular mixtures, despite the addition of equal mass of tissue from each species to a mixture. This is possibly due to species differences in mtDNA copy number per unit mass, and to primer bias, and is particularly marked for fish species (Figure S7).

To assess the performance of VeRIF-ID in mixtures of interest to law enforcement, we analysed a range of seven Traditional East Asian Medicine (TEAM) products (Table S5), from which 203K reads were obtained. About half (100K) remained after filtering, and an average of 9.4K reads per sample remained after barcode demultiplexing. For two samples, no results were obtained, but the species identified within the remaining samples rarely matched those listed on the packaging (when available [Table S5]). Two species of particular interest, the black rhinoceros (*Diceros bicornis*) and saiga antelope (*Saiga tatarica*), were identified within a wax-covered ball intended to tranquillise the nervous system and treat a range of ailments including fever, annoyance, panic, and facial nerve disorder. At the time of manufacture of the TEAM product, both species were listed as critically endangered (www.iucnredlist.org; the saiga antelope’s status was revised in 2023 to near threatened). Packaging in this instance suggested the presence of saiga antelope (as “Cornu Antelopis”), but not black rhinoceros.

### Use of VeRIF-ID for on-site testing

The VeRIF-ID system has been designed to be used on site. We first tested its suitability at Twycross Zoo, UK. We were provided with three samples (a hair, feather, and blood sample) from three known species during a ‘blind’ test. The species of origin was successfully determined in all cases (respectively, Coppery titi, *Callicebus cupreu*s; Demoiselle crane, *Grus virgo*; Francois’s langur, *Trachypithecus francoisi*) in just over 5 h (Table S9).

We next deployed the VeRIF-ID system within a customs zone at Brussels Zaventem Airport, an important gateway for unregulated import of wildlife derivatives, through which around 3.8 tonnes of bushmeat are estimated to enter Europe each month [12]. Bushmeat samples, and fish contaminated with insects (therefore disallowed for import), were seized by customs officials during a passenger baggage compliance action on 24/01/2024. Without disturbance to normal customs activities, the species of origin of these 15 samples was determined in approximately 6 h (Table S10, S11). Samples seized included common livestock (pig [*Sus scrofa*] and cattle [*Bos taurus*]), but also two species of duiker, a small to medium sized antelope. Maxwell’s duiker (*Philantomba maxwellii*) is considered ‘of least concern’ in the International Union for Conservation of Nature (IUCN) classification (www.iucnredlist.org), while bay duiker (*Cephalophus dorsalis*) is listed as ‘near threatened’.

## Discussion

We have described the development of VeRIF-ID (Vertebrate Rapid In-Field Identification via DNA). Our aims were to maximise taxonomic range and the robustness of species identification [71], to reduce time, cost and complexity, and to ensure portability and on-site capability. We chose to use short barcode amplicons to maximise the recovery of data from challenging samples, but clearly there is a balance to be struck between this and the greater certainty of identification offered by longer recovered sequences.

In our test of single-source samples across a wide range of vertebrate classes from lamprey to chimpanzee (Table S3), 91% (75/83) could be assigned to species level, 8% (7/83) to genus, and 1% to tribe level (Figure 2; Table S8). This breadth makes VeRIF-ID a quasi-universal test - a valuable property because it saves the time and expense of specific test development. It is also worth noting that it enables an immediate response, as changes in regulation and market trends alter the species targeted by the IWT [72]. Despite our vertebrate-focused design, we also generated taxonomically useful information for some invertebrate species (Table S5, S10; [66]). As a considerable proportion of illegally traded species consists of invertebrates and plants [73], it would be valuable to multiplex ‘universal’ invertebrate and plant barcode primers in a similar manner, and this should be feasible.

The use of multiple barcodes increases the robustness of identification with respect to barcode drop-out, incomplete or erroneous reference sequence data [32, 33], and the co-amplification or preferential amplification of numts (Figure 2). The 12S and 16S rRNA barcodes were successfully amplified across 100% (83/83) and 99% (82/83) of the species tested respectively, with the 16S sequence missing from only the lamprey, a basal jawless vertebrate. The inclusion of more discriminating COI and CYTB barcodes allowed us to achieve a species-level identification in 20% (15/75) of species where this would not have been possible with the rRNA barcodes alone. In part, this was due to the lack of reference 12S and 16S rRNA sequence data in some species.

Any obtained barcode sequence must be searched against a relevant database of known sequences to effect an identification. Generally, the Barcode of Life Database (BOLD) is preferred since uploaded information must be of a certain quality, and derived from a specimen for which taxonomic assignment can be reviewed [31]. However, BOLD is currently centred around the barcode COI, so not well suited to multi-barcode query. Here we used NCBI’s Nucleotide database, which incorporates GenBank. The latter is a public DNA sequence repository containing non-quality-controlled, unvalidated data in which mistakes are not uncommon [74]. Notably, it has been reported that much of the BOLD data is unauthenticated and itself transferred from GenBank [75]. More generally, any database is susceptible to the misidentification of specimens, a problem for which there is no easy solution [76]. Identification errors are more likely when available data are confined to a single barcode sequence from a single source. By querying multiple different barcode sequences, often derived from different laboratories, the effect of database errors is reduced.

Amplifying numts in addition to, or instead of, target mitogenome sequences can cause problems in species identification. Examples of the mitigation provided by multiple barcodes against numt amplification can be seen in Figure 2. Where numts are amplified and sequenced, they comprise at most two of the four barcodes (prominently in the Felids [34]), while the remaining barcode sequences reflect the endogenous mitogenome.

While we did not attempt any quantitative analysis of mixed multi-species samples, our analysis of lab-prepared mixtures of muscle tissues from up to five different vertebrates shows that VeRIF-ID can generally detect the presence or absence of species reliably. Primer bias leads to dropout of certain barcodes for some species, particularly for fish in the presence of non-fish species. The method is thus susceptible to false negatives and has limited sensitivity, but nonetheless could still detect two critically endangered species in a TEAM sample (Table S5). Such identifications could be followed up with species- or genus-specific primers for confirmation and quantification.

Minimising the time required for species identification is of great practical importance. The first requirement is to ensure that the method can be deployed close to a site of sample seizure. The equipment and reagents required for VeRIF-ID are easily portable by a single operator, and the onboard database obviates the need for an internet connection. A power supply is necessary, and refrigeration is desirable to prolong the life of flow-cells and reagents. Once analysis is commenced, a rapid testing time is important. While we have reduced this considerably, in practice a duration of an hour or less would make testing while a suspect was detained, or a shipment in transit was held, more useful. For VeRIF-ID, the longest step in the process is PCR. Further optimisation of denaturation, annealing and extension time could be explored, and overall time could also be shortened through the use of more efficient thermocyclers than the BentoLabPro, employed here. Such devices include the NextGenPCR™ thermocycler, which has been reported to reduce the time for PCR to around 25 min [77]. However, a compromise is required between speed, expense, and portability - the thermocycler costs about $22K, and weighs 17 kg.

To demonstrate proof of principle, we trialled VeRIF-ID in the customs zone at Brussels Zaventem airport, successfully analysing a variety of seized fish and bushmeat samples from passengers on flights from Addis Ababa, Kigali via Entebbe, and Luanda via Kinshasa. The method allowed 15 samples to be sampled and analysed in around 6 hours, without disruption to normal airport activities (Figure 3, Tables S9, S10). Given the very high traffic of bushmeat via European airports [12, 78], longer term deployment of equipment combined with an efficient workflow is needed to allow higher numbers of samples to be processed during a day. The species identified in Brussels included the bay duiker, which is a reservoir for Ebola virus [79] and has been linked to Ebola spillovers into human populations in West and Central Africa [80]. On-site species identification, potentially followed by pathogen testing in known reservoir species, could provide valuable data for the tracing of pathogens along supply chains and contribute to pandemic prevention preparedness.

**Figure 3:**
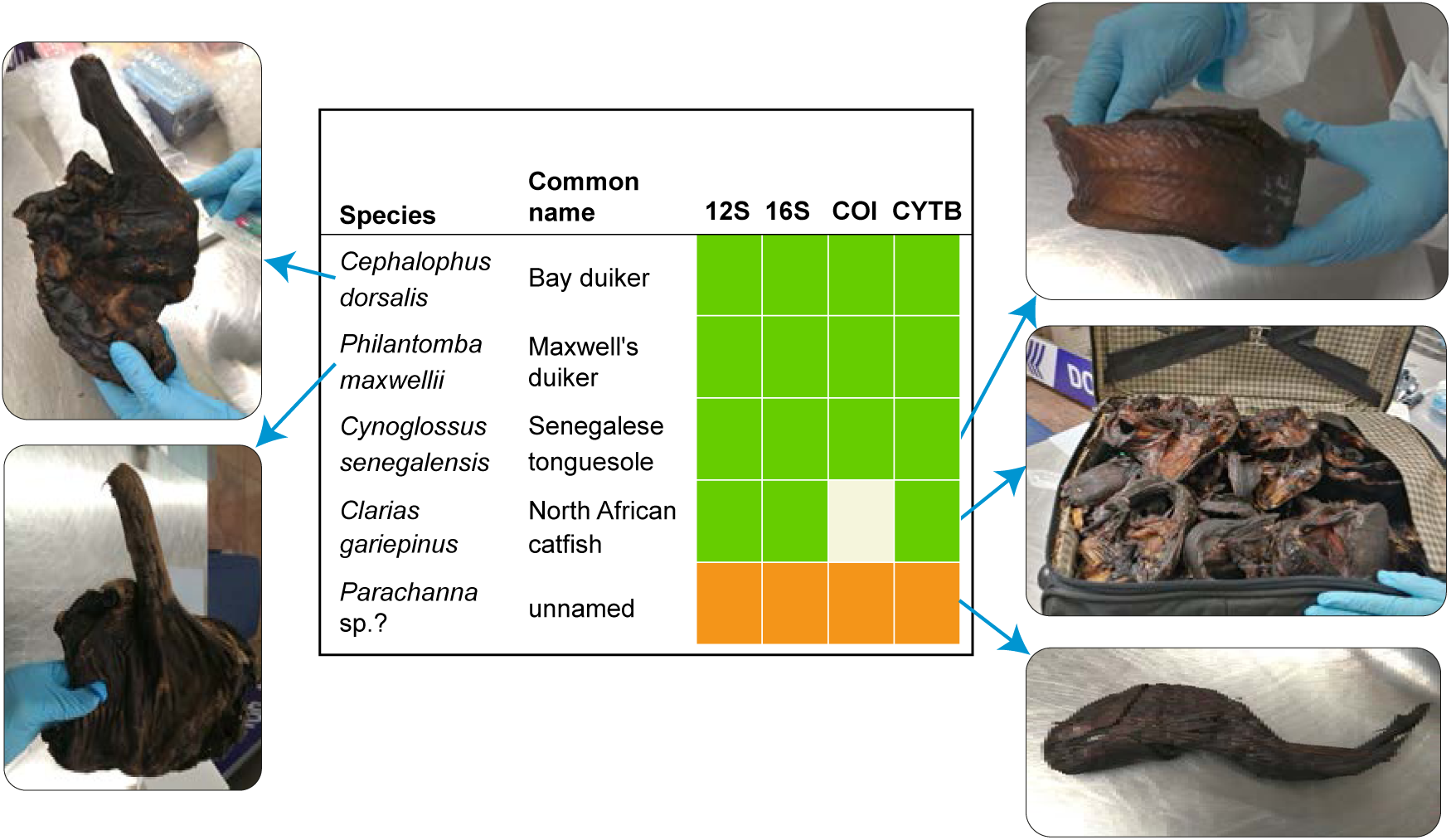
Non-livestock bushmeat and fish species analysed on-site at Brussels Zaventem Airport. Photographs show that bushmeat and fish are dried, and/or smoked. The table gives species and common names (where known) for the five non-livestock samples. Results for the four barcodes are shown to the right (green: amplified and identifying; orange: amplified but no hit in NCBI database (possible undescribed species); white: no amplification). See Table S10 for further information.

Democratising species identification testing demands cheap methods. The total cost of all the required equipment for VeRIF-ID (a MinION sequencer, laptop, BentoLab, centrifuge and a set of pipettes), is relatively low, at approximately $8K. Considering the test itself, while high-throughput laboratory-based DNA barcoding costs as low as $0.10 have been reported [81], in practice forensic testing costs tend to be much higher. Excluding consumables transportation charges, the overall per sample cost of VeRIF-ID, at ∼$43-49 if using a flow cell once (before goods/sales tax, and assuming analysis of 24 samples using a flowcell). This is about 160% cheaper than prices typically charged for species identifications [82]; the on-site nature of the testing also obviates the need for courier charges for sample shipping. Most of our method’s cost is in the sequencing step, which may be reduced by halving reagent volumes during the library preparation step, and either reusing flow cells or through the use of cheaper, single-use Flongle flow cells; see Supplementary Text). Depending on the chosen set-up, per-sample costs can be reduced, for instance if a flow cell is re-used once, the cost per sample decreases to ∼$27-33. Synthesising a set of primer sequences with the indices incorporated within their 5’ tails would allow samples to be pooled immediately after PCR. Only one PCR clean-up would be required, reducing the consumables required, the complexity of the process, and the overall time taken. The slightly cheaper rapid sequencing kit (SQK-RAD114) could then be used.

The simpler a DNA barcoding technique is, the more widely it can be deployed, since relatively little operator expertise is required, and training is simplified. In terms of hands-on abilities, VeRIF-ID is simple, but demands basic lab skills of pipetting; using lyophilised PCR mixes could simplify this.

However, flow cell loading is the most crucial and demanding practical step. Challenges in flow cell loading have been recognised by ONT, which aims to revise the process with a simpler, user-tolerant design. Overall, the most challenging aspect of the method we present here is the need for some familiarity with bioinformatics; developing a simple user interface is therefore an obvious future goal. The development of a user-friendly software with a graphical user interface for use across all commonly used operating systems would greatly increase the accessibility of the workflow. Such software has previously been developed to generate barcodes in real-time [83].

## Declarations

### Ethics approval and consent to participate

This research was approved by the University of Leicester’s Animal Welfare and Ethical Review Body (ref. AWERB/2021/159). All methods were performed in accordance with the relevant guidelines and regulations, and the study is reported in accordance with ARRIVE guidelines (https://arriveguidelines.org). Consent to participate is not relevant.

### Consent for publication

Not applicable.

### Availability of data

Sequence data described in this paper are available from https://doi.org/10.25392/leicester.data.26519668.v1.

### Authors’ contributions

JHW, MAJ conceived and supervised the study; SML collected data; ECP carried out experiments; ECP, JHW carried out data analysis; ECP, JHW, MAJ designed methodology, visualized data and drafted the manuscript. All authors have read, edited, and approved the final manuscript.

## Acknowledgments

We gratefully acknowledge Sarah Bailey, Tara Wilson and Andy Fisher from the Metropolitan Police Service, Lisa Gillespie from Twycross Zoo, and others who contributed DNA samples. We thank Anne-Lise Chaber (University of Adelaide) and Maud Istasse (Belgian Federal Public Service Health) and their colleagues for facilitating testing at Brussels Zaventem airport. This research used the SPECTRE High Performance Computing Facility at the University of Leicester for data analysis.

