## Supplementary Text for "Rapid on-site universal vertebrate species identification via multi-barcode nanopore sequencing"

### 1) Effect on sequence data of using the ONT rapid indexing kit

We chose to use the rapid indexing kit to prepare samples for sequencing given its fast turnaround time and small number of manipulation steps. This kit employs a proprietary ONT transposome complex to fragment double-stranded DNA and attach nanopore indices ('tagmentation') to the resulting 5´ fragment ends (Figure S6; https://store.nanoporetech.com/rapid-barcoding-sequencing-kit-24-v14.html). It is understood [1] that a bacteriophage MuA transposase is used to achieve this. Its action is thought to introduce short (≤100 nt) phage-related DNA segments that may include additional non-viral sequences acquired during previous transposition events [2] to the majority of fragments, as previously noted (e.g. https://github.com/rrwick/Porechop/issues/9, https://community.nanoporetech.com/posts/manufacturing-of-transposa) among users of the kit. This issue is seldom reported in published articles as it has no effect on the kit's primary recommended usage for the analysis of native genomic DNA libraries. Here, the introduction of additional "non-target" bases onto the ends of fragments randomly dispersed within the genome results in their never comprising a significant proportion of contributing base calls anywhere within a consensus sequence. However, when used to sequence targeted PCR products (as in this study), this can be the case at the end furthest from the sequenced primer, because relatively few reads extend to the distal end of each amplicon. Inclusion of 26-nt 5´ extensions on all PCR primers allowed some transposome cuts to be made within the distal primer itself, potentially allowing coverage of every base of the targeted sequence on both strands (addition of primer extensions is now included in the manufacturer's protocols for rapid amplicon sequencing (https://nanoporetech.com/document/rapid-sequencing-v14-amplicon-sequencing-sqk-rbk114-24-or-sqk)).

Given the unpredictable lengths, sequences and positions of these phage-related additions it was not possible to completely remove them bioinformatically from the reads used to generate the forward and reverse consensus sequences, but their impact was largely confined to the end furthest from the sequenced primer site in each case (see Figure S6, where the artefact sequences would appear between the sample-identifying index and the target sequence). These consensus sequences were therefore generally slightly longer than the expected amplicon length, because an insertion into a sequenced fragment can become incorporated into a consensus sequence despite being absent from the majority of reads. Table 1 (below) gives the mean alignment lengths between the top match(es) and the consensus sequences generated. Separate forward and reverse consensus sequences were BLASTed against the NCBI Nucleotide database to detect potential issues arising from numts, which might prime with differential efficiency from each of the forward and reverse primers. Concordance between forward and reverse consensus sequences was required for a species identification and this mitigated issues arising from MuA-associated artefacts, numts and the limited ability of NGSpeciesID to merge short forward and reverse sequence reads.

**Table 1: Mean lengths of consensus sequence alignments.**

| Barcode | Direction | Mean alignment length (bp) | Standard deviation alignment length (bp) | Mean expected insert size based on 843 sequences used for primer design |
|---|---|---|---|---|
| 12S | Forward | 112.5 | 14.2 | 110 (91-117) |
| | Reverse | 114.8 | 13.0 | |
| 16S | Forward | 110.0 | 19.7 | 104 (94 - 111) |
| | Reverse | 113.8 | 16.6 | |
| COI | Forward | 205.9 | 39.3 | 214 |
| | Reverse | 215.9 | 18.8 | |
| CYTB | Forward | 303.6 | 29.7 | 307 |
| | Reverse | 300.8 | 27.8 | |

In all but one case, the additional sequence did not confound accurate species identifications. In the exceptional case, an unusual hybrid 12S rRNA reverse consensus sequence was generated for the bat-eared fox (*Otocyon megalotis*) hair sample sequenced during the single-source flowcell run. This sequence (which has a top BLAST hit for cod) contained the expected 12S rRNA bat-eared fox sequence, a Mu transposase artefact (MuA R1 and R2 recognition sites, and MuA cleavage site), and a 12S rRNA cod sequence generated in the same sequencing batch and possibly attached by residual MuA transposase activity following pooling. Trimming MuA R1 and R2 recognition site sequences [3] using porechop removed the MuA artefact and cod sequences; however, some additional sequences remained at the ends of this 12S rRNA reverse consensus sequence. This artefact was not observed in the 12S rRNA reverse consensus sequence generated with sequencing data called with the newer basecalling software, dorado (v0.8.2) (https://github.com/nanoporetech/dorado) rather than guppy.

As the use of the rapid indexing kit with short amplicon targets is novel, there are no published bioinformatic pipelines to analyse such data. Various modifications to the bioinformatic pipeline (described in the main text) were tested to try and remove these additional variable sequences and obtain amplicon sequences of the expected length. A promising strategy included trimming MuA R1 and R2 recognition sequence sites with porechop after nanopore index demultiplexing. Following MuA sequence trimming, reads were filtered for quality and length with NanoFilt and then assigned to a species ID barcode using porechop by looking for primer sequences (as before). The default trimming function during this second demultiplexing step was disabled, and instead primer trimming was carried out during the consensus sequence generation with NGSpeciesID. While these modifications were largely successful in generating consensus sequences of the expected length, expected barcode sequences generated previously were occasionally not recovered.

An alternative approach to identify the desired amplicon barcode sequence involves the use of EMBOSS megamerger (https://www.bioinformatics.nl/cgi-bin/emboss/megamerger). This tool is used to identify areas of overlap between two sequences, in this case between reverse and forward consensus sequences. If both the forward and reverse consensus sequences generated span the whole amplicon length and do not share any common MuAintroduced sequences at read ends, then the desired amplicon sequence of the expected length can be obtained. As NGSpeciesID does not use nucleotide ambiguity codes during consensus sequence generation, multiple consensus sequences can be generated [4], such that EMBOSS megamerger would need to be used consecutively in these cases.

### 2) Cost per sample

We have described methods to prepare samples for on-site nanopore sequencing. Obtaining amplicons ready for sequencing (DNA extraction, PCR, and PCR clean-up) costs about $4-9.50 per sample across 24 samples depending on whether a KOH- or EDTA-based extraction method is used (see Table 1 below). Approximately 78-33% of this cost is due to the column required for sample clean-up. We used the rapid indexing kit (RBK-114.24) for library preparation. The kit contains the reagents required for six libraries. Using the recommended reagent volumes in the rapid indexing kit leads to a cost of $5.55 per sample for the library preparation step (across a total of 144 samples). It is possible to halve reagent volumes during the adapter dilution and indexing steps to conserve sufficient volumes of these reagents for a further 12 runs. In this case, additional flow cell buffer and flow cell tether are required for these additional runs. These reagents can be purchased separately as sequencing auxiliary vials V14 (EXP-AUX003). Halving volumes where possible, together with the purchase of additional sequencing auxiliary vials can reduce this cost to $2.12 per sample (across 18 libraries for a total of 432 samples). To decrease costs further, 24 samples can be sequenced on the same flow cell more than once. It costs $33.33 per sample if a flow cell is used once, decreasing to $16.66 if used twice, and $11.11 if used three times. It is necessary to wash a flow cell between running different libraries, and the purchase of a flow cell wash kit (EXP-WSH004) would add an additional cost of $0.80 per sample. Alternatively cheaper, single-use Flongle flow cells can also be used instead of a MinION flow cell which would cost $3.19 per sample. Using standard methods, it costs $42.96 - 48.37 per sample from DNA extraction through to sequencing. Halving reagent volumes, purchasing sequencing auxiliary vials, and wash kits would decrease the cost per sample to $23.67-29.08 if a flow cell were washed twice.

**Table 1: Summary of costs under different assumptions.**

Costs are all in US dollars and exclude goods/sales tax.

| **DNA extraction** | **PCR** | **PCR clean-up** | **Library preparation** | | | **Sequencing** | | | | Per sample total cost (across 24 samples) |
|---|---|---|---|---|---|---|---|---|---|---|
| | | | Rapid indexing kit (RBK-114.24; $800 for 6 reactions) | Rapid indexing kit (RBK-114.24; volumes halved) + Sequencing auxiliary vials (EXP-AUX003; $115 for 12 reactions) | Rapid indexing kit (RBK-114.24; volumes halved) | Flow cell ($800) - used once | Flow cell + Flow cell wash kit (EXP-WSH004; $115 for 6 reactions)- used twice | Flow cell + Flow cell wash kit (EXP-WSH004; $115 for 6 reactions)- used thrice | Flongle flow cell ($918 for 12) | |
| <0.10 - 5.51 | 0.88 | 3.10 | 5.55 | | | 33.33 | | | | 42.96-48.37 |
| | | | 5.55 | | | | 17.47 | | | 27.10-32.51 |
| | | | 5.55 | | | | | 11.91 | | 21.54-26.95 |
| | | | | 2.12 | | 33.33 | | | | 39.53-44.94 |
| | | | | 2.12 | | | 17.47 | | | 23.67-29.08 |
| | | | | 2.12 | | | | 11.91 | | 18.11-23.52 |
| | | | | | 2.78 | | | | 3.19 | 10.05-15.46 |

### 3) VeRIF-ID protocol

**List of equipment & reagents:**

- Pipettes (P2, P20, P200, P1000) and tips
- 1.5-mL LoBind DNA Eppendorf tubes
- 1.5-mL Eppendorf tubes
- 0.2-mL PCR tubes
- Disposable biopsy punches, scissors, or scalpel
- KOH-based lysis buffer and Tris-based neutralisation buffer
- Bento Lab Pro
- Table-top centrifuge with 12 places
- Adapters for 0.2-mL tubes
- Type-It Microsatellite PCR kit
- Primer mixes
- Molecular biology grade water
- MinElute PCR purification kit (containing Buffer PB, Buffer PE, and spin columns)
- Rapid indexing ('barcoding') kit (RBK-114.24)
- MinION device
- Laptop
- MinION flow cell
- Timer
- Standard PPE (gloves, bench covers, etc.)

**A) DNA extraction using an alkaline lysis buffer**

1. Obtain 1-2 $mm^3$ piece of claw, tissue, shed skin, feather calamus, spawn, quill, 3 µL of blood, or 5 – 10 hairs.
2. Place sample in a 0.2-mL PCR tube.
3. Add 100 µL lysis solution (200 mM KOH, 2 mM EDTA, 0.2% (v/v) Triton X-100).
4. Heat on the BentoLab Pro for 10 mins at 98°C. Spin down.
5. Add 300 µL of 100 mM Tris-HCl to a 1.5-mL Eppendorf tube.
6. Transfer the solution in the 0.2-mL PCR tube to the 1.5-mL Eppendorf tube containing the Tris-HCl solution.
7. Flick or invert to mix and spin down.

**B) PCR amplification**

1. Prepare PCR: 6.25 µL Type-It Master Mix, 5.03 µL of water, 0.22 µL of primer mix (for a final concentration of 0.3 µM of CYTB primers, 0.25 µM of COI and 16S rRNA primers, and 0.08 µM of 12S rRNA primers), and 1 µL of DNA extract in a 0.2-mL PCR tube.
2. Mix by flicking and spin down.
3. Place reaction(s) on the BentoLab Pro PCR machine and heat at 95°C for 5 min, then 5 cycles at 95°C for 30 s, 45°C for 90 s, 72°C for 30 s, 35 cycles at 95°C for 10 s, 45°C for 10 s, 72°C for 40 s, and a final extension at 68°C for 2 min.
4. Spin down.

**C) PCR clean-up**

1. Add 62.5 µL of MinElute Buffer PB to the 0.2-mL PCR tube and mix by pipetting.
2. Transfer solution to a MinElute spin column.
3. Centrifuge at maximum speed for 1 min.
4. Add 750 µL of MinElute Buffer PE to the column.
5. Centrifuge at maximum speed for 1 min.
6. Discard flow-through and place the column back in its collection tube.
7. Centrifuge at maximum speed for 1 min.
8. Transfer column to a new 1.5-mL Eppendorf tube.
9. Add 10 µL of water to the column.
10. Wait 1 min.
11. Centrifuge at maximum speed for 1 min.
12. Transfer 5 µL of purified PCR product to a new 0.2-mL PCR tube.

**D) Library preparation**

1. Spin down indexes and adapter solutions and mix by pipetting.
2. Add 0.5 µL of rapid index (1-24) to 5 µL PCR product in the 0.2-mL PCR tube. Mix by flicking and spin down.
3. Heat on the BentoLab Pro for 2 mins at 30°C for 2 mins, then 80°C for 2 mins. Spin down.
4. Pool 4 µL of each sample in a 1.5-mL LoBind Eppendorf tube. Mix by flicking and spin down.
5. Dilute the rapid adapter by adding 0.75 µL of the rapid adapter to 1.75 µL of the adapter buffer in a 1.5-mL LoBind Eppendorf tube. Pipette to mix.
6. In a 1.5-mL LoBind Eppendorf tube, add 11 µL of the pooled indexed samples with 1 µL of the diluted adapter. Mix by flicking and spin down.
7. Incubate at room temperature for 5 mins.

**E) Sequencing and data analysis**

(for a detailed protocol see: https://community.nanoporetech.com/docs/prepare/library_prep_protocols/rapid-sequencing-v14-amplicon-sequencing-sqk-rbk114-24-or-sqk/v/raa_9198_v114_revh_29nov2023/priming-and-loading-the-spoton-flow-cell?devices=minion)

1. Thaw and spin down sequencing buffer (SB), library beads (LIB), flow cell flush (FCF), flow cell tether (FCT).
2. Place flow cell in MinION device.
3. Start MinKNOW and start flow cell check.
4. Add 30 µL of FCT directly to a single-use FCF tube or 1,170 µL FCF in a new 1.5-mL Eppendorf tube. Mix by pipetting.
5. Slide the priming port clockwise on the flow cell.
6. Insert a P1000 pipette (set to 200 µL) into the priming port and turn the wheel until a small amount of buffer enters the pipette tip.
7. Add 800 µL of flush buffer (prepared in step 4) to the flow cell through the priming port.
8. Wait for 5 mins.
9. In a new 1.5-mL LoBind Eppendorf tube add 37.5 µL of SB, 25.5 µL of LIB (mixed by pipetting to resuspend beads prior to addition), and 12 µL of library (prepared in step D7).
10. Lift the SpotON sample port cover.

11. Add 200 µL of flush buffer (prepared in step 4) via the priming port.
12. Pipette to mix library prior to loading. Add 75 µL to the SpotON sample port one drop at a time.
13. Close the SpotON sample port cover and the priming port.
14. Start sequencing with MinKNOW. During the sequencing set-up specify the library preparation kit used (RBK-114.24), a minimum read length of 20 bp, enable barcode trimming, ensure read filtering is on, and specify that a high accuracy basecalling model is used (assuming a sufficiently powerful computer/laptop is used. For details of recommended specifications see: https://community.nanoporetech.com/requirements_documents/minion-it-reqs.pdf)Obtain at least 1K passed reads across most indexed samples.
15. Stop sequencing.
16. Use NanoFilt to select for reads between 100 and 500 bp.
17. Use Porechop to assign reads to species ID barcodes.
18. Use NGSpeciesID to generate consensus sequences for reads assigned to a species ID barcode.
19. Use the consensus sequences as a BLAST query against a local NCBI nucleotide database. Specify the output file format using the options "7 qseqid qlen evalue qcovs pident staxids stitle scomnames".
20. For each sequence's BLAST result, filter to retain a unique result for each taxonomic identifier. Further filter the results to retain only the top hit(s).
21. Optionally, steps 16-20 can be implemented as a simple script.
