## Supplementary Figures for "Rapid on-site universal vertebrate species identification via multi-barcode nanopore sequencing"

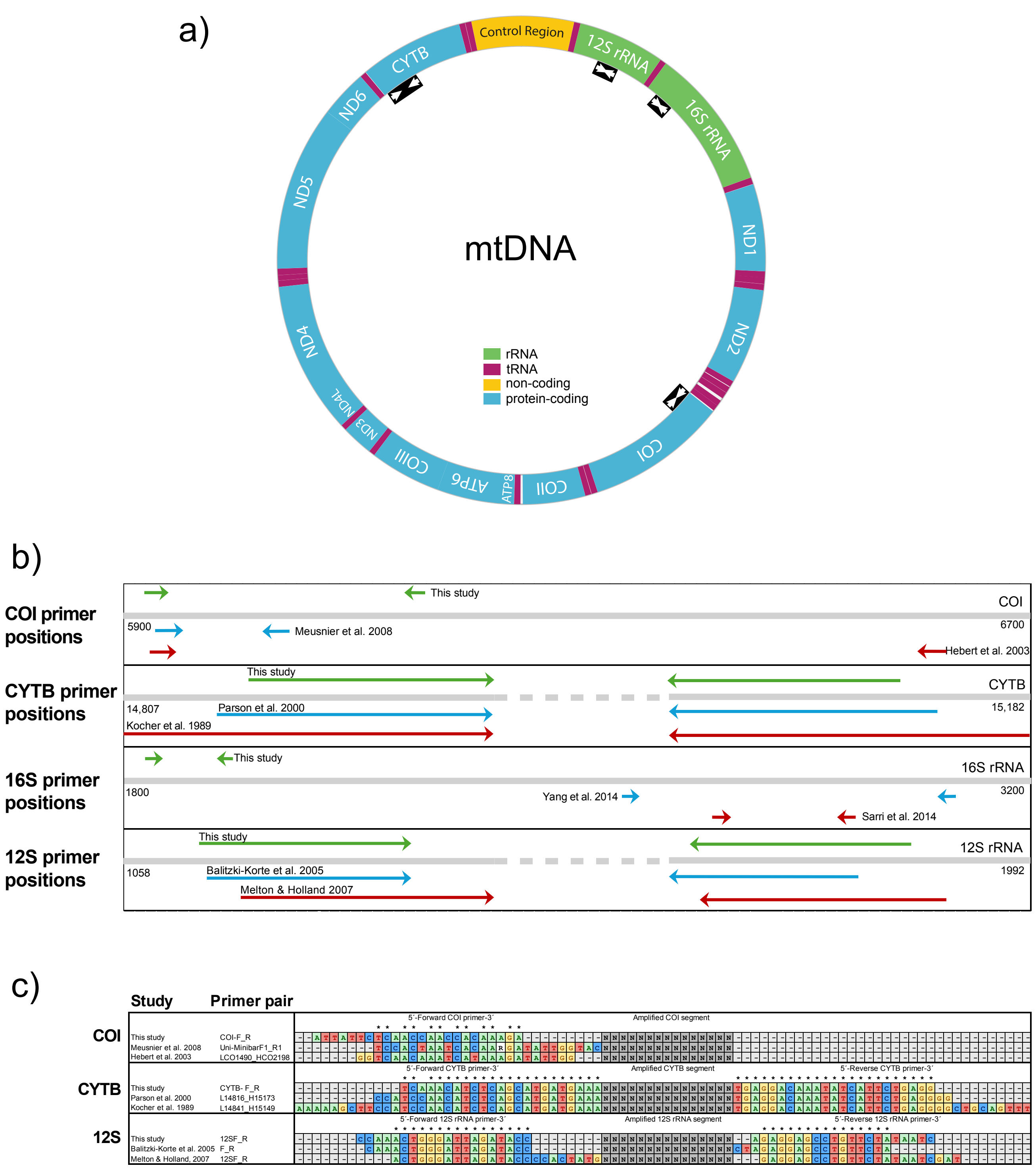


**Figure S1: Barcode primer positions in mtDNA, and relationships to previously designed primers.**

a) Schematic diagram of typical vertebrate mtDNA, with approximate positions of the four barcode sequences. Primer sequences are given in Table S2.
b) Schematic representation of the relationship between the primers in this study and previously published primers. Numbers indicate coordinates, based on the human mtDNA reference sequence (NC_012920).
c) Detailed comparison of primer sequences for barcodes COI, CYTB, and 12S rRNA. 16S rRNA primers are not shown since they have no overlaps with previously described primers.

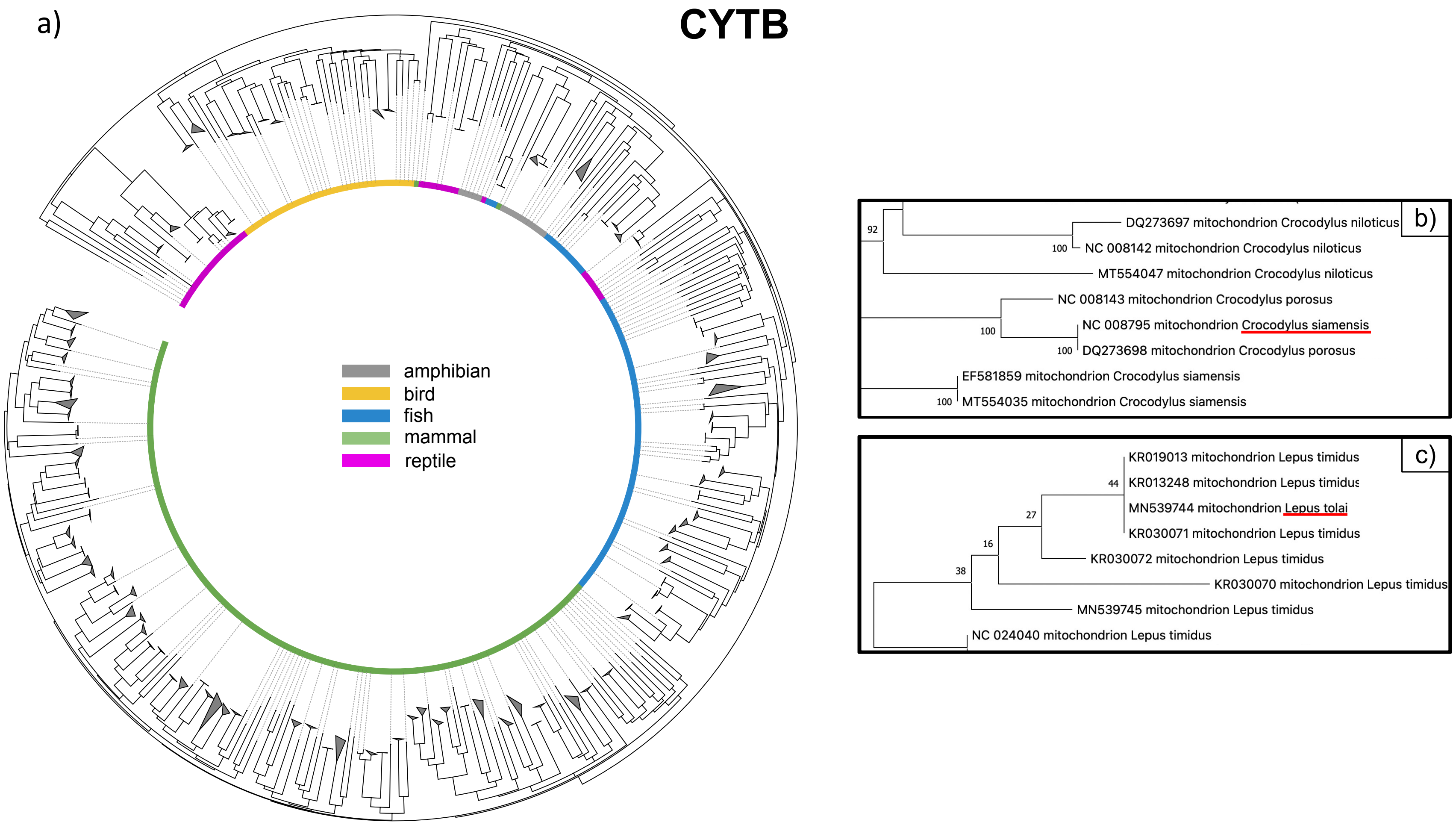


**Figure S2: Phylogenetic tree based on cytochrome b.**

a) A NJ-K2P tree with 100 bootstrap replicates was calculated using MEGA (Hall, 2013) and displayed using the interactive Tree Of Life (Letunic and Bork, 2021) for the chosen 307-bp segment of cytochrome b. Clades have been collapsed when multiple sequences of the same species (or closely related species) were present. Instances where species differentiation was not possible are displayed in panels b) and c). Inconsistent species identifications are highlighted by red underlining.

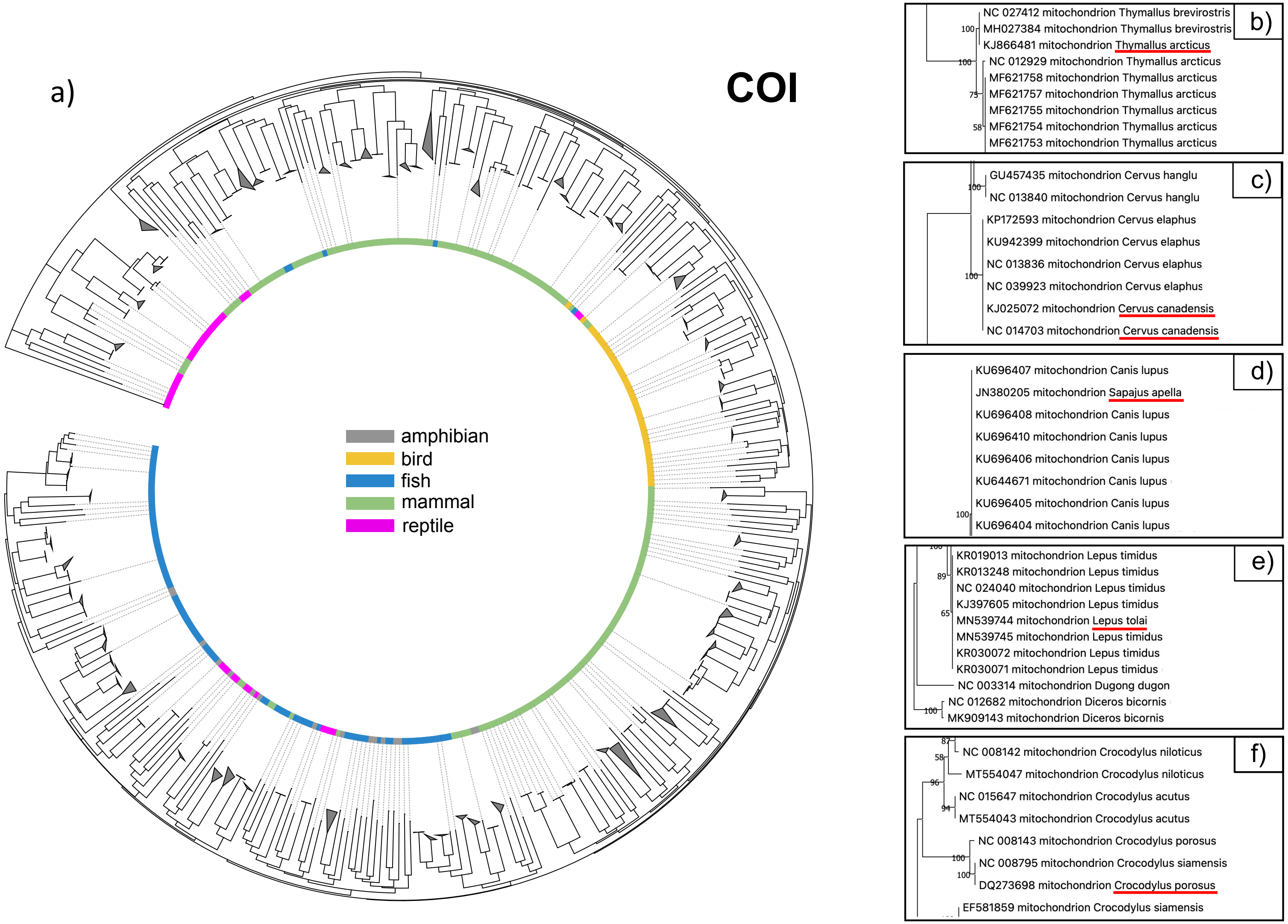


**Figure S3: Phylogenetic tree based on cytochrome c oxidase subunit I.**

a) A NJ-K2P tree with 100 bootstrap replicates was calculated using MEGA (Hall, 2013) and displayed using the interactive Tree Of Life (Letunic and Bork, 2021) for the chosen 214-bp segment of cytochrome c oxidase subunit I. Clades have been collapsed when multiple sequences of the same species (or closely related species) were present. Instances where species differentiation was not possible are displayed in panels b) to f). Inconsistent species identifications are highlighted by red underlining.

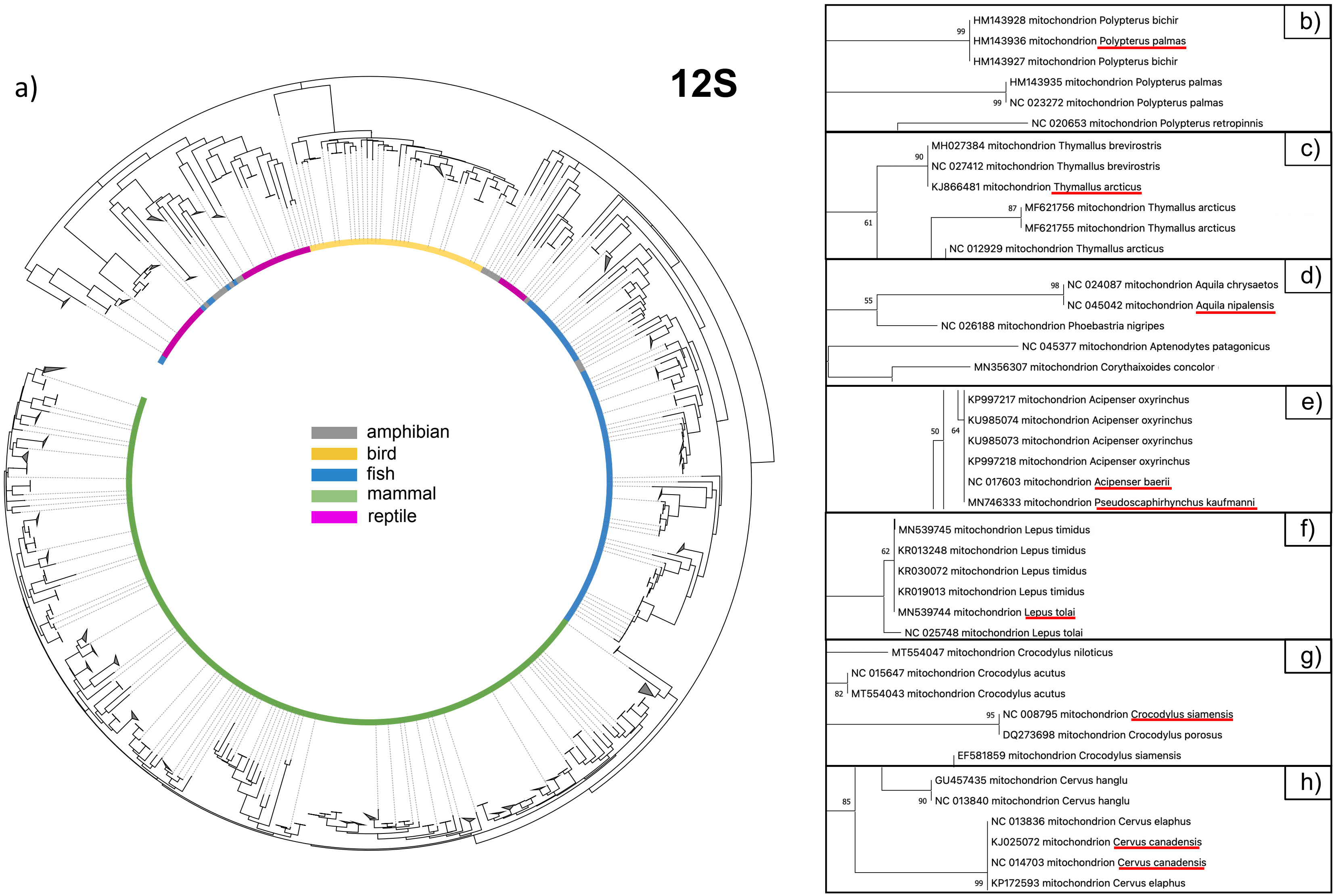


**Figure S4: Phylogenetic tree based on 12S ribosomal RNA.**

a) A NJ-K2P tree with 100 bootstrap replicates was calculated using MEGA (Hall, 2013) and displayed using the interactive Tree Of Life (Letunic and Bork, 2021) for the chosen 111-bp segment of 12S rRNA. Clades have been collapsed when multiple sequences of the same species (or closely related species) were present. Instances where species differentiation was not possible are displayed in panels b) to h). Inconsistent species identifications are highlighted by red underlining.

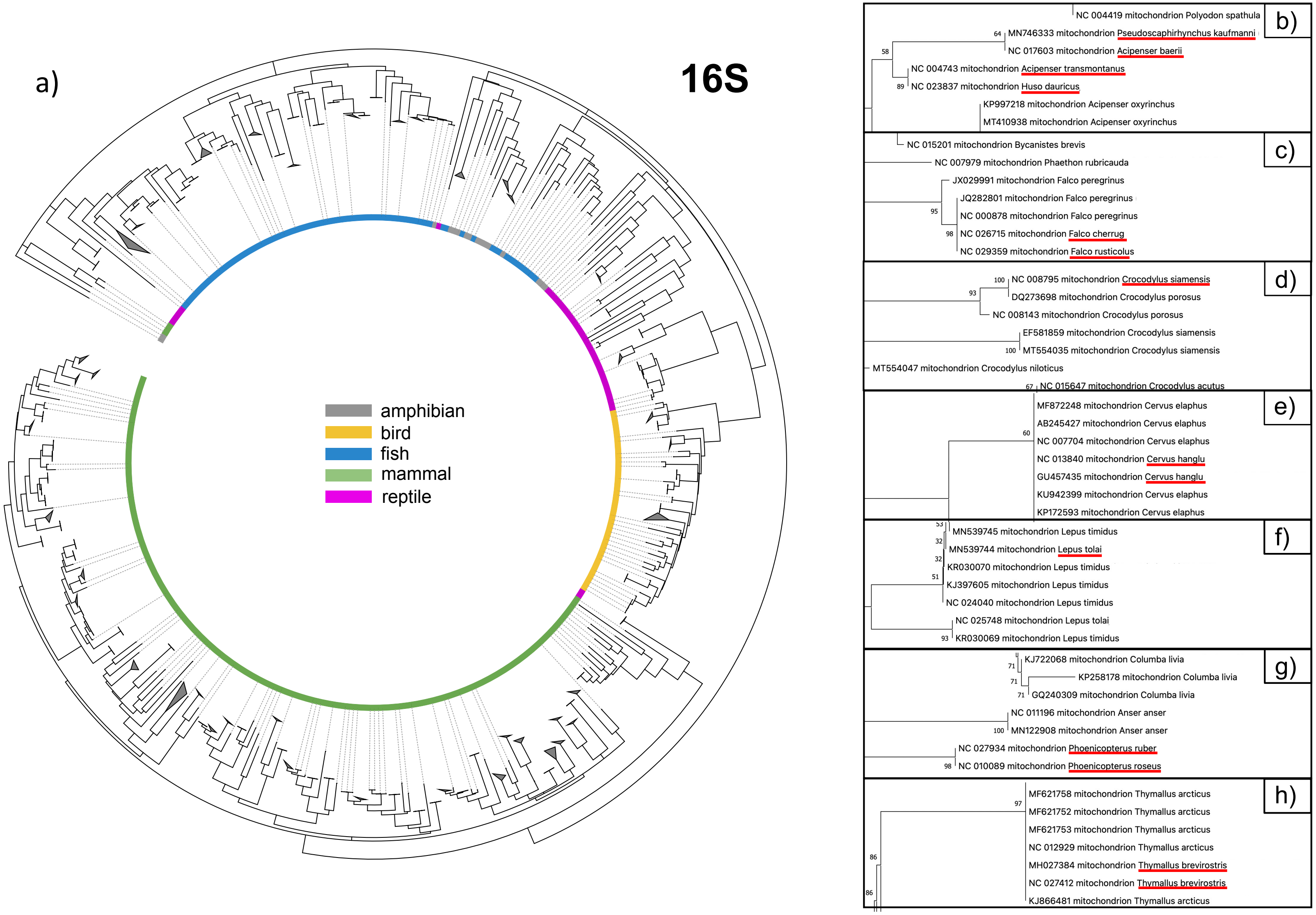


**Figure S5: Phylogenetic tree based on 16S ribosomal RNA.**

a) A NJ-K2P tree with 100 bootstrap replicates was calculated using MEGA (Hall, 2013) and displayed using the interactive Tree Of Life (Letunic and Bork, 2021) for the chosen 103-bp segment of 16S rRNA. Clades have been collapsed when multiple sequences of the same species (or closely related species) were present. Instances where species differentiation was not possible are displayed in panels b) to h). Inconsistent species identifications are highlighted by red underlining.

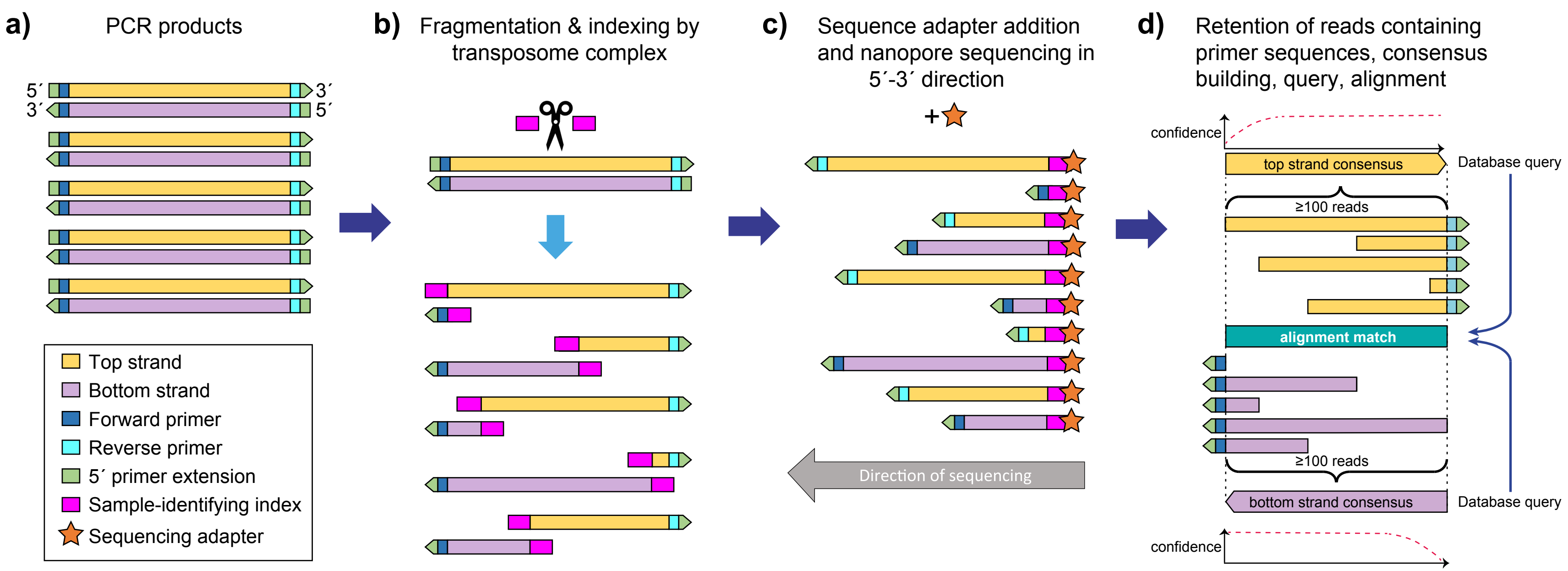


**Figure S6: Action of the rapid indexing kit and generation of sequences for database query.**

**a)** PCR products are generated by primers with 26-nt 5´ extensions to increase the target size for transposome complex binding. **b)** Transposome complex introduces a double-stand break at a random point within each duplex and adds a sample-identifying index to each of the two internal free 5´ ends. **c)** Following sequencing adapter addition, sequencing proceeds from each adapter through the nanopore in the 5´-3´ direction. **d)** In bioinformatic processing, a read is only retained if it contains a primer sequence at the distal end (primer sequences are then trimmed - not shown), and reads are used as input with NGSpeciesID for consensus sequence generation. This leads to reduced confidence at the proximal end of each strand consensus. Top- and bottom-strand consensuses are used separately for database query and only results with consensus sequences based on at least 100 independent reads and generated in both directions are considered (though see main text).

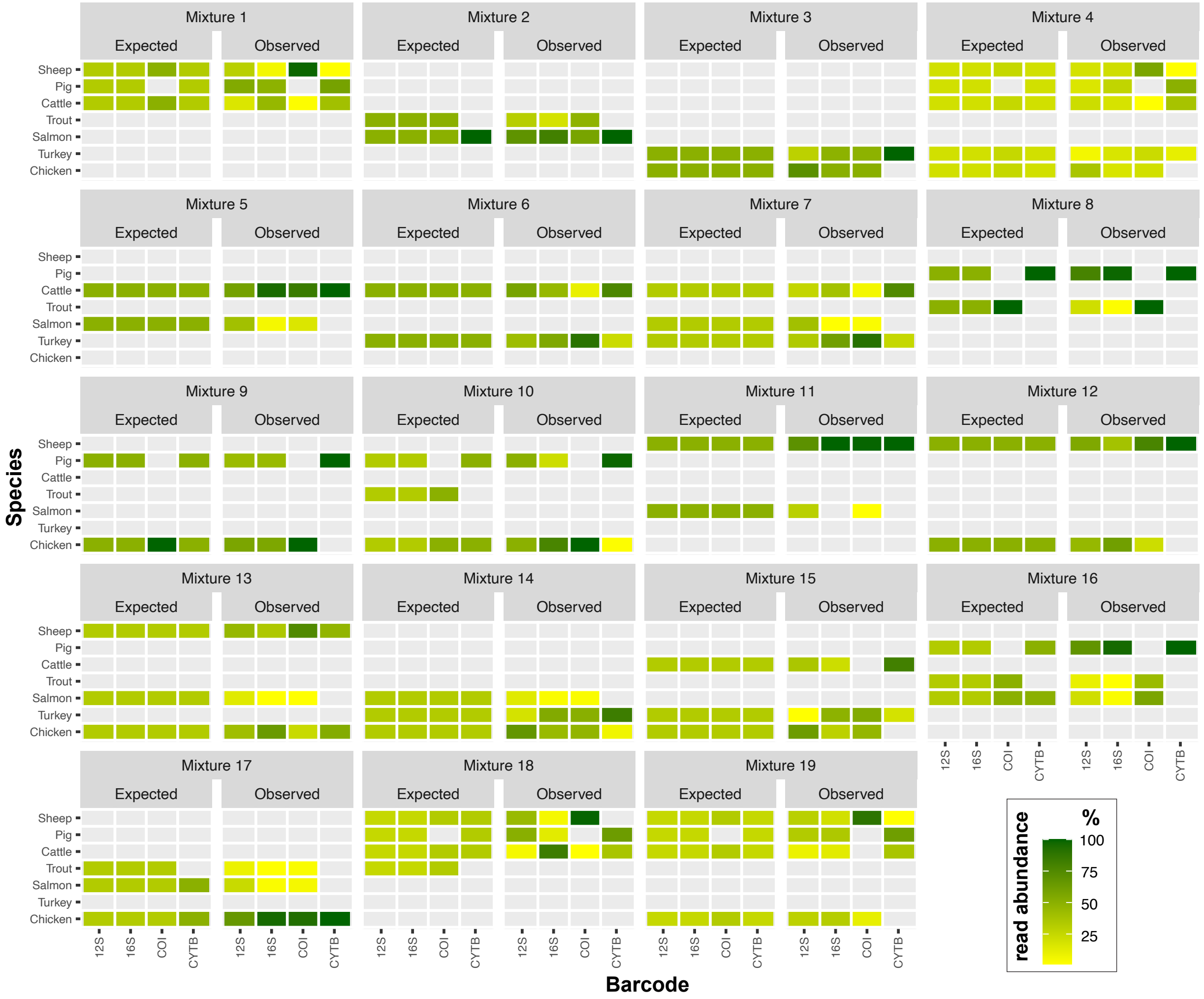


**Figure S7: Heatmap displaying the relative percentage of reads assigned to a given species within 19 lab-prepared mixtures.**

For each mixture, the first column indicates the relative proportion of reads expected for each species present in the mixture for that indicated barcode in the minibarcode tetraplex. The second column for each mixture indicates the relative proportion of reads assigned to the barcode of each of the detected species based on the highest number of supporting reads of that species' barcode consensus sequence. In sequence processing consensus sequences with >20 supporting reads were retained. A species was only considered present if forward and reverse consensus sequences were generated for at least two species ID barcodes.
